# Estradiol Action at the Median Preoptic Nucleus is Necessary and Sufficient for Sleep Suppression in Female Rats

**DOI:** 10.1101/2020.07.29.223669

**Authors:** Philip C. Smith, Danielle M. Cusmano, Shaun S. Viechweg, Michael D. Schwartz, Jessica A. Mong

**Affiliations:** Department of Pharmacology, University of Maryland Baltimore, Baltimore Md

## Abstract

To further our understanding of how gonadal steroids impact sleep biology, we sought to address the mechanism by which proestrus levels of cycling ovarian steroids, particularly estradiol (E2), suppress sleep in female rats. We showed that steroid replacement of proestrus levels of E2 to ovariectomized female rats, suppressed sleep to similar levels as those reported by endogenous ovarian hormones. We further showed that this suppression is due to the high levels of E2 alone, and that progesterone did not have a significant impact on sleep behavior. We found that E2 action within the Median Preoptic Nucleus (MnPO), which contains estrogen receptors (ERs), is necessary for this effect; antagonism of ERs in the MnPO attenuated the E2-mediated suppression of both non-Rapid Eye Movement (NREM) and Rapid Eye Movement (REM) sleep. Finally, we found E2 action at the MnPO is also sufficient for sleep suppression, as direct infusion of E2 into the MnPO suppressed sleep. Based on our findings, we predict proestrus levels of E2 alone, acting at the MnPO, mediate sex-hormone driven suppression of sleep in female rats.

**Competing Interest Statement:** MDS is now an employee of Jazz Pharmaceuticals; the views expressed in this manuscript are solely the authors’ and do not reflect the views, policies or procedures of Jazz Pharmaceuticals.

## Introduction

Primary sleep disorders are among the most common medical conditions. Compared to men and boys, women and girls are twice as likely to experience sleep disruptions and insomnia throughout their lifespan.^1^ This discrepancy may be due to a multitude of factors, including chromosomal sex differences and psychosocial factors such as anxiety prevalence. However, clinical data suggest the increased risk of sleep disruption in women emerges at puberty and has been associated with fluctuations in ovarian steroids, particularly estrogens. This evidence suggests that gonadal steroids and biological sex are significant risk factors for sleep disruptions.^2^ Apart from the clinical finding that sex steroids may affect sleep behavior and architecture, the mechanisms and site of action underlying a link between steroids and sleep disturbances remain a gap in our knowledge. In particular, despite a growing understanding of sleep regulatory mechanisms, how and where estrogens influence the sleep circuitry is poorly understood. To gain a better understanding of sleep disruptions and insomnia, we must first understand how and where estrogens influence basic sleep mechanisms in females.

Historically, the majority of sleep studies have been performed in men or male animals,^2^ a deeply unfortunate occurrence that has neglected the impact female animal models can have on illuminating ties between ovarian steroids and sleep. The paucity of basic studies investigating sex differences in sleep has resulted in an unclear picture on the nature of those differences. Sleep patterns in female rats are exquisitely sensitive to the natural fluctuations of ovarian steroids.^3–4^ Multiple studies in rats show that during proestrus, when estrogen and progesterone are elevated, sleep time is significantly reduced compared with other phases of the estrous cycle.^5–6^ Exogenous hormone replacement is observed to recapitulate this phenotype^7^ in both rats and mice. In these studies, estradiol predominately suppresses dark phase sleep and has little or no effect in the light phase. Thus, a key paradigm for studies of hormonal modulation of sleep has been the use of hormone replacement in ovariectomized rodents,^2^ which can provide hormonal stability that bypasses the rapid hormonal changes inherent to the 4-day estrous cycle in rats.

Several nuclei are thought to stimulate sleep, with two nuclei of the preoptic hypothalamus of key importance. The Ventral Lateral Preoptic (VLPO) and Median Preoptic (MnPO) are thought to be key originators of this pathway.^2^, ^8–9^ Numerous studies have shown neurons in the VLPO and the MnPO are involved in sleep-regulatory mechanisms.^10–13^ Both the VLPO and MnPO (i) have a predominant number of sleep-active cells (i.e., the number of Fos-ir neurons increase following episodes of sleep but not waking)^9, 14^ (ii) have a high concentration of neurons with elevated discharge during sleep compared to waking (i.e. sleep-active discharge pattern),^15^ and (iii) are thought to promote and sustain sleep by inhibiting arousal centers.^11–13, 16^ A feed-forward loop has been identified in which the MnPO both inhibits the wake system of the lateral hypothalamus and stimulates the VLPO, which itself serves as an inhibitor of the wake nuclei.^9^ As the MnPO and VLPO serve as upstream modulators of wake systems, direct manipulations of these nuclei can therefore be employed to test the subsequent sleep-wake pathway.

Steroid receptors are present throughout the brain and prevalent on multiple sleep-regulating nuclei such as the hypothalamus^17^ and basal forebrain.^18^ Previous work in rodents implicates the VLPO in particular as a key site of mediating E2 actions over sleep.

In adult ovariectomized females, E2 decreases activation of sleep-active VLPO neurons^19^ and downregulates levels of lipocalin-type prostaglandin D synthase (L-PGDS), the enzyme responsible for the production of prostaglandin D2 that potently promotes sleep.^20–21^ Estrogens also decrease expression of wake-inhibiting adenosine 2A receptors, suggesting a potential alternate mechanism for an inhibitory impact on homeostatic sleep pressure.^22^ However, these findings are complicated by studies showing that the VLPO is not a major site of estrogen sensitivity.^23^ Instead, we hypothesize that it is the MnPO which may be most responsible for mediating E2 action over homeostatic sleep pressure. Blocking E2 action directly in the median preoptic nucleus of female rats attenuates E2 suppression of sleep.^19^

In the current study, we sought to expand our knowledge of the mechanisms through which ovarian steroids regulate sleep-wake behavior in adult female rats and specifically determine if the preoptic area is the locus of these effects. Using an exogenous hormone replacement model that mimics the estrous cycle levels and timing of E2 and progesterone, we first tested whether E2 alone is sufficient to induce changes in sleep-wake behavior and sleep homeostasis or if progesterone has additional actions. Second, using pharmacological manipulations of local estrogenic signaling in the preoptic area sleep nuclei, with ER antagonism and local E2 infusion, we investigated whether E2 is necessary and sufficient to induce changes in sleep-wake behavior and homeostasis. These experiments test the tripartite hypothesis that E2 acting (1) alone at the MnPO is (2) necessary and (3) sufficient to induce the changes in sleep seen with high hormone levels in cycling female rodents.

## Materials and Methods

### 1. Animals

Adult female Sprague–Dawley rats (250-350g) were purchased from Charles River Laboratories (Kingston, N.Y.) and housed in the Laboratory Animal Facilities at the University of Maryland, School of Medicine under a reversed 12 h: 12 h dark: light cycle (Lights on at 9PM for Chapter II and experiments 1 and 3 in Chapter III, lights on at 6AM for Chapter III, experiment 2.) with food and water available ad libitum. In all experiments, zeitgeber time 0 (ZT 0) represents Lights ON. Due to logistical limitations, all experimental procedures were run in multiple cohorts, with all experimental groups represented in each cohort. All procedures were performed in accordance with the National Institutes of Health guide for care and use of laboratory animals. All experiments were approved by and were in accordance with the guidelines of the University of Maryland Institutional Animal Care and Use Committee.

### 2. Gonadectomies and Transmitter/Cannula Implantation

All surgeries were conducted under isoflurane anesthesia. All animals were ovariectomized (OVX) or castrated according to standard protocol and simultaneously implanted with TL11M2-F40-EET transmitters (Data Sciences International, St. Paul, Minn.). Briefly, animals were OVX using a dorsal incision followed by isolation and removal of the ovaries bilaterally. Using the OVX incision for females, a bipotential-lead transmitter (DSI Inc., St. Paul, Minn.) was implanted intraperitoneally. Another incision was made along the midline of the head and neck to expose the skull and neck muscle. Two burr holes were drilled asymmetrically and stainless-steel screws (Plastics One, Roanoke, Va.) were implanted at 2 mm anterior/1.5 mm lateral and 7 mm posterior/1.5 mm lateral relative to the bregma. The four transmitter leads were threaded subcutaneously to the head incision. Electroencephalographic (EEG) leads were wrapped around the screws and secured with dental cement. Electromyogram (EMG) leads were implanted directly in the dorsal cervical neck muscle, approximately 1.0 mm apart, and sutured in place.

For experiments that required local injections into preoptic area nuclei, guide cannula were implanted. Three types of guide cannula were used. For MnPO infusion, a single guide cannula (C315G, 26-gauge; Plastics One) targeted to the MnPO was implanted at a 9° angle at the stereotaxic coordinates 0.45mm posterior/ +1.0mm lateral/ 6.5mm ventral relative to bregma. For VLPO infusion, a bilateral guide cannula (C235G, 26-gauge; Plastics One) targeted to the VLPO was implanted at the stereotaxic coordinates 0.1mm posterior/ 1.0mm lateral/ 7.0mm ventral relative to bregma. In all cases, the cannula and EEG leads were secured together with dental cement. Upon insertion of the cannula, the opening was closed with a matching dummy provided by the respective cannula manufacturer. The skin along the head was sutured around the guide and dummy cap, and the dorsal incision was closed with wound clips. Fig. 11 is a representative image of the guide cannula placement. All animals were treated with antibiotic ointment and topical lidocaine as well as carprofen (5 mg/kg) postoperatively and then allowed 7 days to recover before the start of the experiments.

### 3. Data Collection and Sleep Scoring

Home cages with the telemeter-implanted animals were placed on receiver bases that continuously collected EEG and EMG data at 500Hz and transferred the data to a PC running Ponemah Software (DSI Inc, St. Paul, Minn.). Digitized signal data was scored off line using NeuroScore DSI v3.3.9 (DSI Inc, St. Paul, Minn.). The EEG/EMG signals were parsed into 10 second epochs. A Fast Fourier transform (Hamming window, 4096 samples) was used to break down the EEG frequency bands (Delta (0.5-4-Hz), Theta (4-8Hz), Alpha (8-12Hz), Sigma (12-16Hz), Beta (16-24Hz) Gamma (24-50Hz) and Total (0.5-50Hz)). The mean of the absolute value was calculated for the EMG data (bandpass 20-200Hz). These data were exported to Matlab (Matlab R2015, Mathworks, Natick, Mass.) where vigilant states were automatically scored using a custom program developed by the lab which has been shown to be ~88% in agreement with hand scored traces.^24^

### 4. EEG Spectral Analysis

To further test whether estradiol influences sleep homeostasis, NREM Slow Wave Activity (NREM-SWA; a marker of sleep homeostasis), was assessed via EEG spectral distributions of NREM sleep bouts. From Neuroscore, the EEG power spectra were calculated for slow wave activity (0.5-4Hz), and into 0.25Hz stepwise bins between 0.5-20Hz. Each power bin was normalized to the mean total power from the 24-hour baseline recording. Slow wave activity was averaged into 1h bins during the light phase. Spectral power was then averaged into 6-hour epochs (ZT times 0-6, 6-12, 12-18, and 18-0). For both slow wave activity and spectral power, artifact was removed if data were 3 standard deviations from the mean of the surrounding 24 data points. The algorithm is described in more detail in Smith et al 2021.^24^

### 5. Steroid Treatments

For experiment 1, all animals were administered 50uL of sesame oil on Day 1. Animals were subsequently administered 5 μg 17-β-estradiol benzoate in 50uL sesame oil (E2; Sigma-Aldrich, St. Louis, MO) on Day 2, and 10 μg E2 in 100uL sesame oil 24 h later on Day 3, or equivalent amounts (50uL/100uL) of sesame oil vehicle, through subcutaneous flank injections. On Day 4, animals received a dose of 500mg progesterone in 50μL sesame oil vehicle, or sesame oil vehicle control. Experiment 2 follows the same timing paradigm with the omission of the Day 4 progesterone injection. For experiment 3, 5ug cyclodextrin-encapsulated E2 in 5uL saline (Sigma-Aldrich), or 5uL free cyclodextrin vehicle (Sigma-Aldrich), was infused directly into the MnPO in each of three successive injections 24 hours apart. Experimental manipulation and sleep data collection was performed at times from 4 to 36 hours after the second hormonal injection (see specific experiments below).

### 6. Drugs and Infusion Paradigm

Animals in experiment 1 comprised a single group that all received ovariectomy and identical hormone replacement as described above, with each animal’s individual baseline serving as a control. Animals in experiment 2 were randomly assigned into either the vehicle (VEH; 0.25% dimethyl sulfoxide (DMSO) in sterile saline) or ICI (50ng in 0.25% DMSO in sterile saline; Sigma-Aldrich) infusion groups, and reversed the following week for a second round of infusions. For targeted infusions to the VLPO, the dummy stylet was removed and replaced with a 33-gauge micro-infusion needle, which extends 2.0mm below the tip of the guide cannula. For targeted infusions to the MnPO, the dummy stylet was removed and replaced with a 33-gauge micro-infusion needle (Plastics One), which extends 0.5mm below the tip of the guide cannula. The needle was connected to a Hamilton 1705 RNR 50ul syringe (Hamilton, Reno, NV) via polyethylene tubing. A BASi Bee pump and Bee Hive controller (Bioanalytical Systems, Inc., West Lafayette, IN) was used to deliver ICI or VEH at a rate of 0.1μl/min. Following infusion, the needle remained in place for 5 minutes to ensure diffusion. ICI or VEH was infused 3 times per injection: (i) 6-12h prior to, (ii) 30 minutes prior to and (iii) 12h after injections (Fig. 10). Similarly, animals in experiment 3 received targeted infusions to the MnPO of cyclodextrin-encapsulated E2 or cyclodextrin vehicle; the dummy stylet was removed and replaced with a 33-gauge micro-infusion needle (Plastics One), which extends 0.5mm below the tip of the guide cannula. The needle was connected to a syringe and controller as described above. The setup was used to deliver cyclodextrin-encapsulated E2 or cyclodextrin vehicle at a rate of 0.1μl/min. Following infusion, the needle remained in place for 5 minutes to ensure diffusion.

### 7. Cannula Placement Verification

At the end of each experiment, animals were overdosed with a ketamine/ acepromazine mix before being transcardially perfused with 0.9% saline + 2% sodium nitrite followed by 4% paraformaldehyde in 0.05M KPBS. The brains were removed and post-fixed overnight in 4% paraformaldehyde. Brains were cryoprotected in 30% sucrose in KPBS, frozen on dry ice, and stored at −80°C. Each brain was cut on a cryostat along the coronal plane at 30μm thick into 4 series and stored in an ethylene glycol-based storage solution at −20°C. Sections in each series are separated by 120μm.

Sections corresponding to the VLPO and MnPO from one series were mounted on 2% gelatin-coated slides. The slides were processed for cresyl violet (0.1% solution; cresyl violet acetate, Sigma-Aldrich) staining to examine cannula placement. VLPO hits were counted as placement within sections 32-36 of the brain atlas^25^ and MnPO hits were counted as placement within sections 33-35. For experiment 2 in the VLPO, one animal was a miss and excluded from the study, while 3 animals were euthanized prior to completion and removed from the study. For all MnPO cannulations, animals with cannula placement outside of this area were removed from analysis; 3 animals were removed. There was 1 animal in experiment 2 whose cannula placement was a miss but remained in the analysis; this animal was infused with VEH and her behavior was not different from hits.

### 8. Statistical Analysis

All data are represented as mean ± SEM. Two-way, repeated measure ANOVAs followed by Sidak post-hoc tests were run for each vigilance state to determine if direct VLPO and MnPO infusions significantly altered E2 effects on sleep-wake. Since this was a within-animal study, systemic injection (oil vs. E2) was the repeated factor and infusion (VEH vs. ICI) was the independent factor. An *a priori* comparison of interest was between the VEH and ICI infused E2 days of analysis. We ran an unpaired t-test to compare means on the E2 day between VEH and ICI infused animals. T-tests were used to compare E2 and VEH MnPO infusions and two-way, repeated measure ANOVAs followed by Sidak post-hoc tests were run for analysis across the phase in 1h bins. Mann-Whitney U nonparametric tests were run to analyze differences between mean percent changes of each vigilance state. All statistical tests were conducted using the Graph Pad Prism program (San Diego, CA) on a PC. In all figures (*) denotes significance at p<.05, (**) denotes significance at p<.01, (***) denotes significance at p<.001, and (****) denotes significance at p<.0001.

## Results

### Experiment 1. Estradiol is the Ovarian Steroid that Predominantly Influences Sleep-Wake Behavior in the Adult Female Rat

Our laboratory and others have demonstrated that exogenous E2 administration, which mimics the levels and timing of the fluctuations in endogenous hormones, markedly reduces time spent in NREM and REM sleep with a concomitant increase in the time spent in wake.^1, 6–7, 19, 26–27^ However, these previous findings have only analyzed the 24 hours after the last injection of E2. To further explore and establish this model of ovariectomy followed by hormone replacement, which mimics the natural rise of E2 to peak proestrus level^80^ and also recapitulates the sleep patterns of intact females,^19^ we recorded and analyzed sleep across the treatment paradigm. We ovariectomized (OVX) female rodents and replaced E2 and progesterone globally through subcutaneous injection, in a cycle formulated to mimic endogenous hormone steroid levels. This replacement paradigm consists of an oil dose on Day 1 designed to mimic metestrus, a low 5ug E2 dose on Day 2 designed to mimic diestrus, and a high 10ug E2 dose on Day 3 designed to mimic proestrus.^6^ (See “Steroid Treatments” in methods.) The advantage of this established model (ovariectomy; OVX + exogenous E2 replacement that mimics the gradual natural rise of E2 to peak proestrus levels) is the standardization and reproducibility of circulating E2 levels on specific recording days. Following the second E2 treatment, the animals were divided into two groups and administered a physiological dose of progesterone (1mg; based on our established findings^19^ or vehicle (referred to as Post E2) on Day 4. (Fig. 1).

**Fig. 1.**
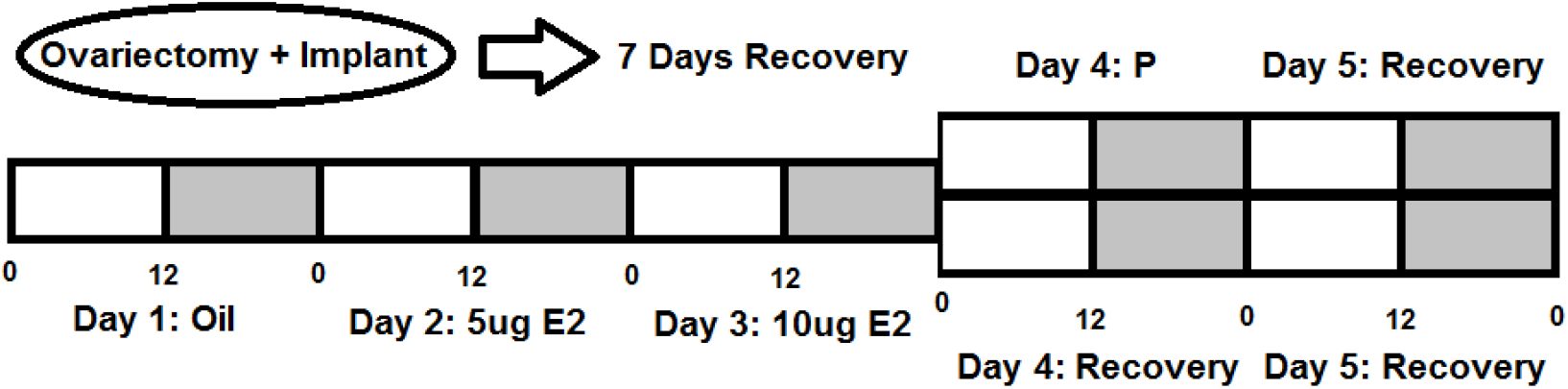
Timeline of Hormonal Recapitulation Experiment. Ovariectomized Female Sprague-Dawley Rats (n=17) were administered an oil injection on Day 1, 5ug of E2 on Day 2, and 10ug of E2 on Day 3. On Day 4, the animals were split into two groups, with one group being administered progesterone to mimic the estrous hormone milieu and one group being administered vehicle to solely examine solely the effect of E2. Sleep times were measured using EEG/EMG telemetry (DSI Inc. St. Paul, Minn.)

#### a. Proestrus-Level Estradiol is Sufficient to Suppress Sleep

As anticipated, in the dark phase, E2 significantly increased the time spent in wake at the expense of NREM. This change was present on the day of high E2 administration. (Fig. 2–3–4) compared to the oil baseline.

**Fig. 2.**
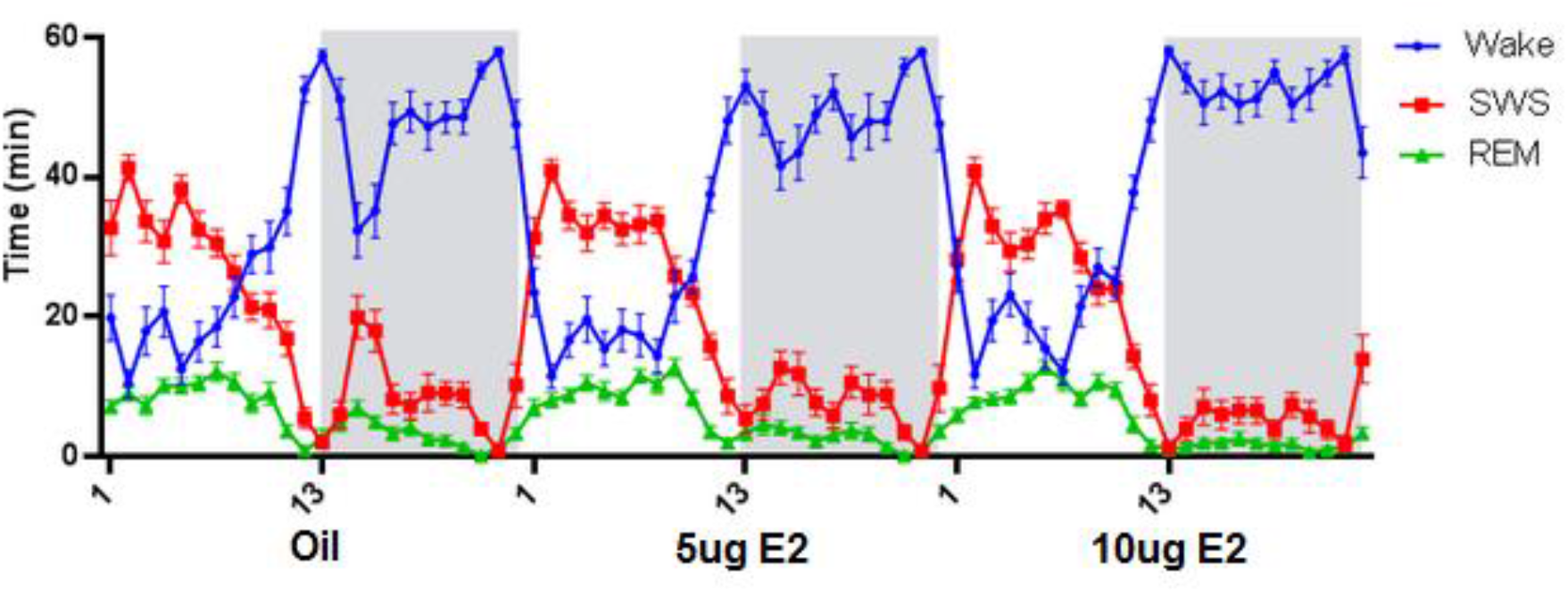
Proestrus-level E2 Suppresses Sleep and Increases Wake. Using a within animal design, we recorded EEG/EMG data from OVX adult rats treated with our standard dosing paradigm of 2 injections of E2 24 hours apart. On the day of high-dose E2 administration (Day 3), mimicking proestrus hormone levels, there is an increase in wake time and decrease in slow wave sleep time. This sleep change mimics the change in sleep on proestrus in naturally cycling rodents. (Repeated measure ANOVA; Wake main effect of treatment: F(2,29)=13.37, p<0.0001); (Repeated measure ANOVA; NREM main effect of treatment: F(2,29)=14.15, p<0.0001).

Conversely to the changes in wake (Fig. 3A), there was a significant decrease in NREM sleep time in the dark phase on the day of high E2 administration (Fig. 3B). E2 administration also significantly decreased REM sleep. (Fig. 3C)

**Fig. 3.**
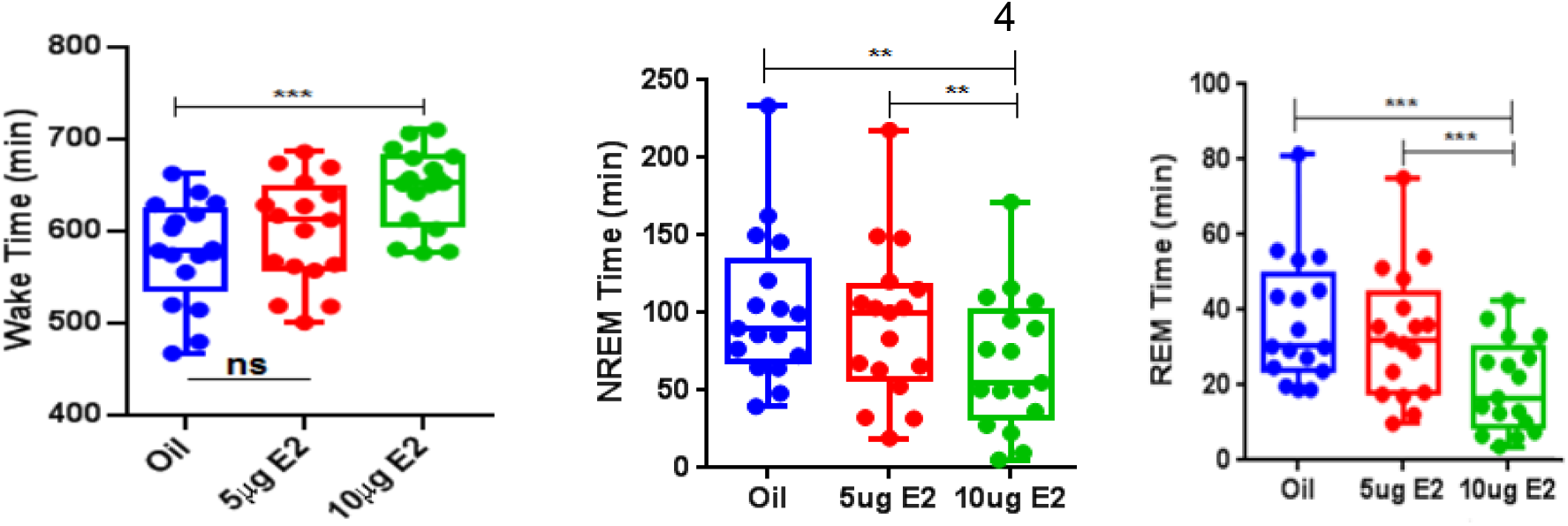
A-B-C. E2 Increases Total Dark Phase Wake Time and Decreases Total NREM and REM Sleep Time. **(A)** In the dark phase, animals showed a significant increase in wake time relative to oil on the day of high-dose E2 treatment (p=.0005). The low-dose E2 treatment did not have a significant effect on wake, while the progesterone-treated animals did not show a significant difference in wake from their E2-only treated counterparts on the same day. (Repeated measure ANOVA; Wake main effect of treatment: F(2,29)=13.37, p<0.0001). **(B)** The high-dose E2 showed a significant decrease in NREM sleep time in the dark phase compared to both oil (p<.01) and low dose E2 (p<.01). (Repeated measure ANOVA; NREM main effect of treatment: F(2,29)=14.15, p<0.0001) **(C)** The high-dose E2 showed a significant decrease in REM sleep time in the dark phase compared to both oil (p<.001) and low dose E2 (p<.001). (Repeated measure ANOVA; REM main effect of treatment: F(2,29)=15.43, p<0.0001)

It is interesting to note that on the day analogous to proestrus (Post E2), E2 treatment abolished the mid-phase siesta by markedly increasing wake at ZT 14-16 compared to the baseline and low E2 days (Fig. 4). This effect was not present in the light phase, with no significant change in light phase sleep time noted across any of the treatment days. (Fig. 5)

**Fig. 4.**
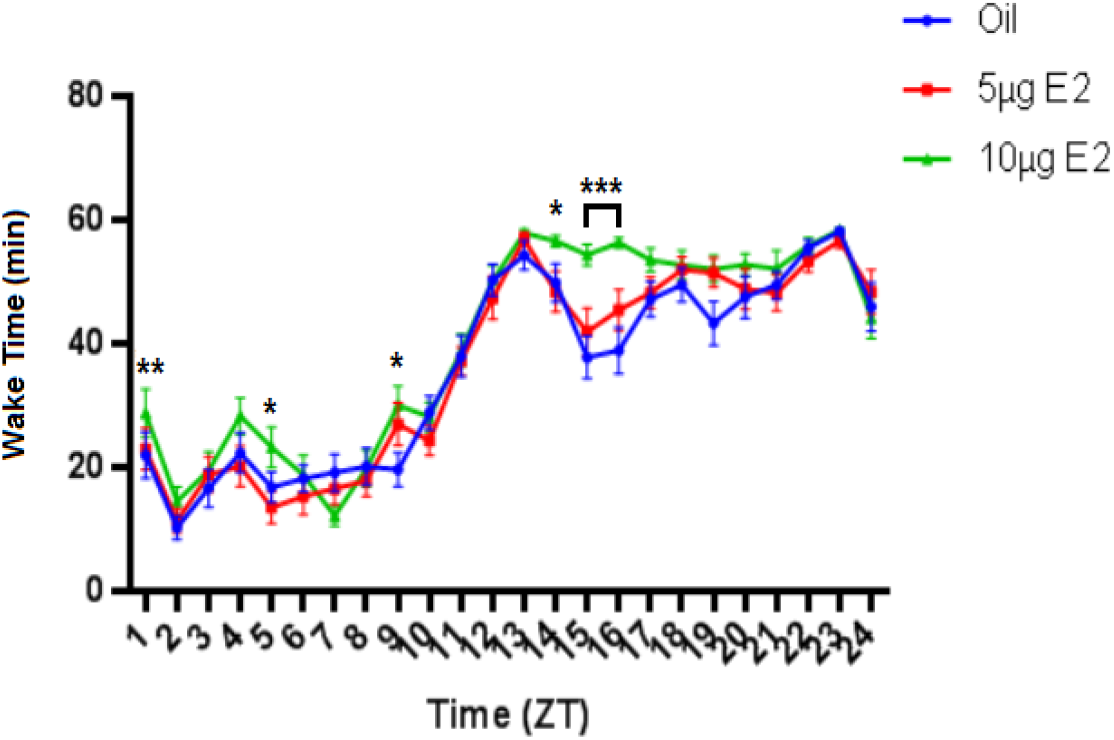
E2-mediated Sleep Suppression is Most Prevalent in the Early Dark Phase. Analysis of the sleep time by hour across the treatment days, shows that wake time is not significantly different between the oil and low dose E2 at any treatment time. However, the high-dose E2 treatment showed a significantly higher wake time at six hourly time points, with the effect being particularly pronounced in the early dark phase (ZT-14-16) (Repeated Measure 2-way ANOVA; Main effect of treatment F(2,48)=46.50; p<0.0001. Sidak’s multiple comparison test, Oil vs. 10ug E2, ZT 1 p<.01, ZT5 p<.05, ZT 9 p<.05, ZT 14 p<.05, ZT 15 p<.001, ZT 16 p<.001)

#### b. Progesterone Has No Significant Additional Effect on Sleep-Wake States

After showing the effects of E2 alone, a question remained over whether progesterone, which also rises on the afternoon of proestrus in natural cycling females, is influencing sleep and wake. Thus, to further validate our model, following the second E2 treatment, on Day 4 the animals were divided into two groups and administered a physiological dose of progesterone (P; 1mg, which is a dose relevant to endogenous proestrus levels)^28^ or vehicle (referred to as Post E2). Moreover, we also analyzed sleep times with and without progesterone. We found that progesterone had no significant effect on sleep-wake states, either wake, NREM, or REM, when compared to the analogous Day 4 (Post E2) day (Fig. 6A-B-C).

**Fig. 5.**
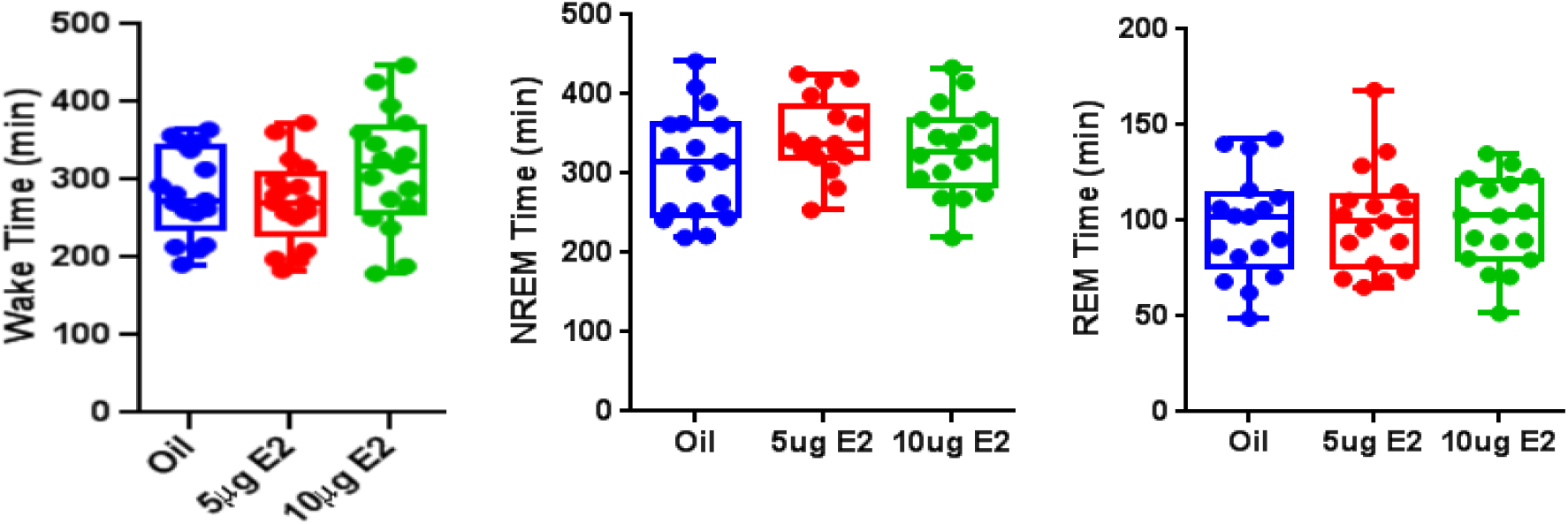
A-B-C. E2 does not Affect Total Sleep or Wake Time in the Light Phase. There is no significant difference in **(A)** wake time, **(B)** NREM sleep duration, or **(C)** REM Sleep Duration across any treatment.

**Fig. 6.**
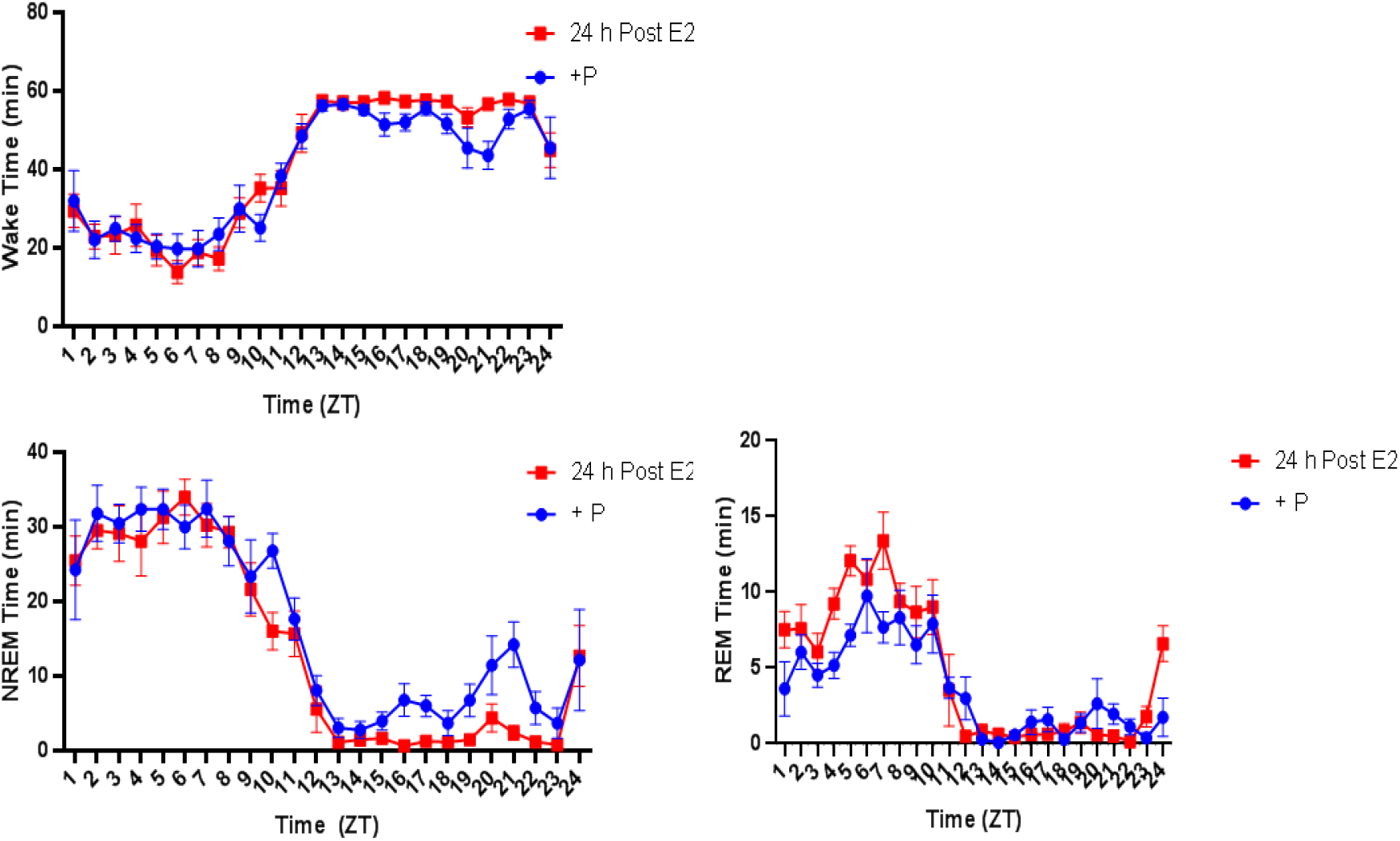
A-B-C. Progesterone does not Affect Sleep Times. To compare for the effect of Progesterone, we split the cohort into two treatment groups on Day 4, one receiving progesterone on the day after high estrogen, recapitulating the estrus phase, and one receiving no additional treatment. There was no significant difference between the groups in **(A)** wake time, **(B)** NREM sleep time, or **(C)** REM Sleep Time. (Mixed Effect Model: treatment x time, Wake: F(4.08,46.9)=1.12, NREM: F(4.08,46.9)= 1.15, REM F(4.08,46.9)=1.18)

Furthermore, we also analyzed NREM delta (0-4 Hz) power through Fourier transformation of the EEG signal, a widely used^29–30^ measure of the depth of homeostatic sleep, both with and without progesterone. When normalizing delta power to each animal’s baseline oil day (Day 1), we found no significant change in the relative delta power difference between progesterone-treated and untreated animals. However, in both groups, there was a decrease relative to oil baseline in the E2-treated animals (Fig. 7). Thus, these findings validate the ovariectomy + exogenous E2 (alone) model as a reliable experimental system that is amenable to local manipulation of sleep-active nuclei to more directly test how estrogens modulate the sleep-circuits and elicit changes in sleep behavior, and show that global E2 action alone is sufficient to recapitulate sleep changes in naturally cycling rodents.

**Fig. 7.**
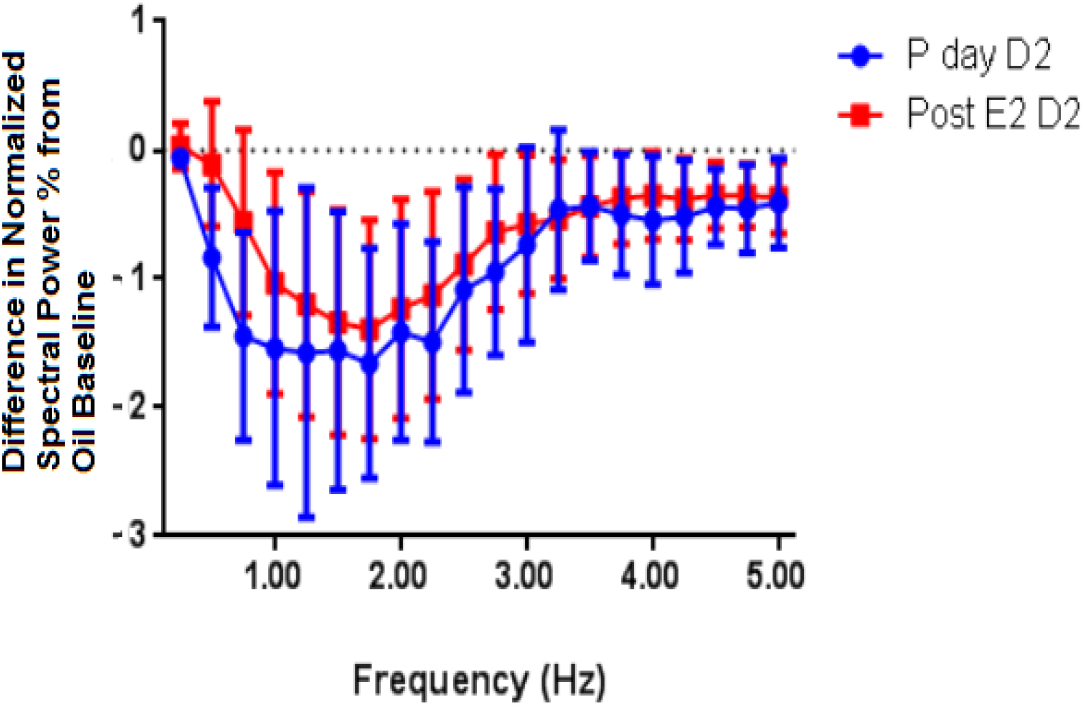
Progesterone Produces No Change in EEG Spectral Power. We compared power spectra of the progesterone and non-progesterone treated groups on Day 4 relative to their own Day 1 oil baseline. In both groups, we see a decrease in spectral power on the day of progesterone treatment, with no significant difference between the progesterone treated and E2 only groups.

#### c. Estradiol Decreases NREM-SWA Spectral Power

To investigate these EEG power spectrum findings (Fig. 7) further, we examined the differences in EEG power between oil-treated and post-E2/E2+P animals, comparing each animal’s Day 1 and Day 4 readings. The power frequency distribution of dark phase NREM-SWA from the females used in the progesterone experiment was compared between Oil versus the day post-E2, which represents the period of the greatest NREM sleep loss following E2 administration, with or without progesterone. We found that there was no significant change in power across all sleep states during either the first half (Fig. 8A) or second half (Fig. 8B) of the light phase. However, during the dark phase, significant changes were observed. Given the significant decrease in dark phase NREM sleep (~40%), we expected the NREM-SWA frequency distribution in the 0.5-4.0 Hz bands to be significantly greater following E2 treatment. However, E2-treated animals, both with and without progesterone, showed a significant *decrease* in EEG power in the low frequency ranges, particularly in the delta band. (Fig. 8C) Comparisons of the normalized percent of total power spectral distribution revealed that E2 significantly decreased the dark phase power of the 1-2.5 Hz bands, which typically represent the highest level of cortical synchronization and thus high-quality sleep, suggesting a decreased level of deep, homeostatically restorative sleep. In the second half of the light phase, the effect of lowered EEG power with E2 was less-pronounced, (Fig. 8D) but appeared to be present over a broader range of frequencies, including in the theta band. These results suggest that the decrease in NREM sleep time in E2 also manifests as a decreased level of deep, homeostatically restorative sleep, and that E2 may attenuate the build-up of SWA under normal physiological conditions.

**Fig. 8.**
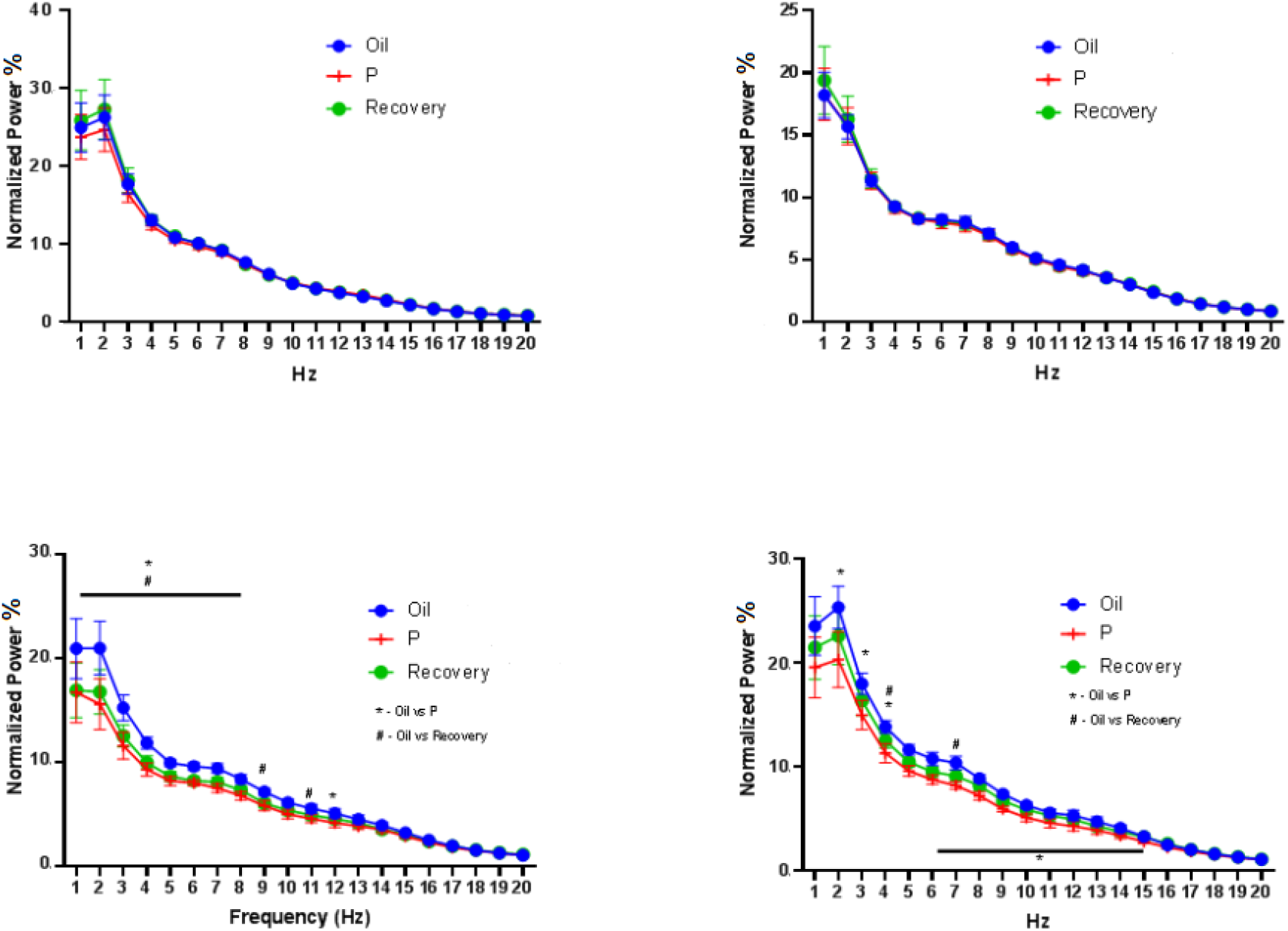
A-B-C-D. E2 Decreases Spectral Power in the Dark Phase, Particularly in the Delta Band. We examined the spectral power across all animals on Day 1, the day of oil treatment (Oil), Day 4, the day of progesterone treatment (P) following E2, or the following Day 5 (Recovery). (See Fig. 1) There was no significant change in spectral power at any frequency in the light phase, either the ZT 0-6 early portion **(A)** or the ZT 6-12 later portion **(B)**. In the dark phase, however, there was a pronounced decrease in normalized power at low frequencies, particularly in the delta and theta ranges in the first half of the dark phase ZT 12-18 **(C)**, when sleep times are most affected by E2. These low power ranges have been shown to be important for homeostatically restorative sleep. (REML Mixed-Effects model with multiple comparisons, main effect of hormone, F (19, 200) = 105.9, p<.001, interaction of hormone X time, F (57, 580) = 1.484, p<.05) In the second half of the dark phase ZT18-0 **(D)** there was a significant decrease in the Theta and Alpha frequency ranges as well as the Delta. (REML Mixed-Effects model with multiple comparisons, main effect of hormone, F (19, 200) = 141.0, p<.0001, interaction of hormone X time, F (57, 580) = 1.901, p<.001)

### Experiment 2. Estrogen Receptor Antagonist Action at the MnPO but not the VLPO of Adult OVX Females Attenuates Estradiol Mediated Suppression of Sleep

#### a. ER Alpha Expression is Present at High Levels in the Female MnPO but not the VLPO

Building on those results, we next attempted to determine if the POA circuitry is necessary to drive these effects. Over the past decade, numerous studies using various techniques have convincingly demonstrated that neurons in the VLPO and the MnPO are involved in sleep-regulatory mechanisms.^8–9^ The VLPO and MnPO reportedly have complementary roles in the maintenance of sleep, as they both (i) have a predominant number of sleep-active cells (i.e., the number of Fos-ir neurons increases following episodes of sustained sleep but not sustained waking),^9, 14^ (ii) have a high concentration of neurons with elevated discharge rates during both NREM and REM sleep compared to waking (i.e. sleep-active discharge pattern),^15^ and (iii) are thought to function to promote and sustain sleep by inhibiting key arousal centers via descending GABAergic (MnPO) and GABAergic/galaninergic (VLPO) projections.^11–13, 16^ Here we investigate whether estrogen receptors were present in these sleep-associated nuclei. Immunocytochemistry using polyclonal antibodies against ER alpha demonstrated a significantly greater population of ER alpha positive cells in the MnPO compared to the VLPO (Fig. 9). These findings show that the MnPO appears to be the major seat of E2 sensitivity in these active sleep circuits.

**Fig. 9.**
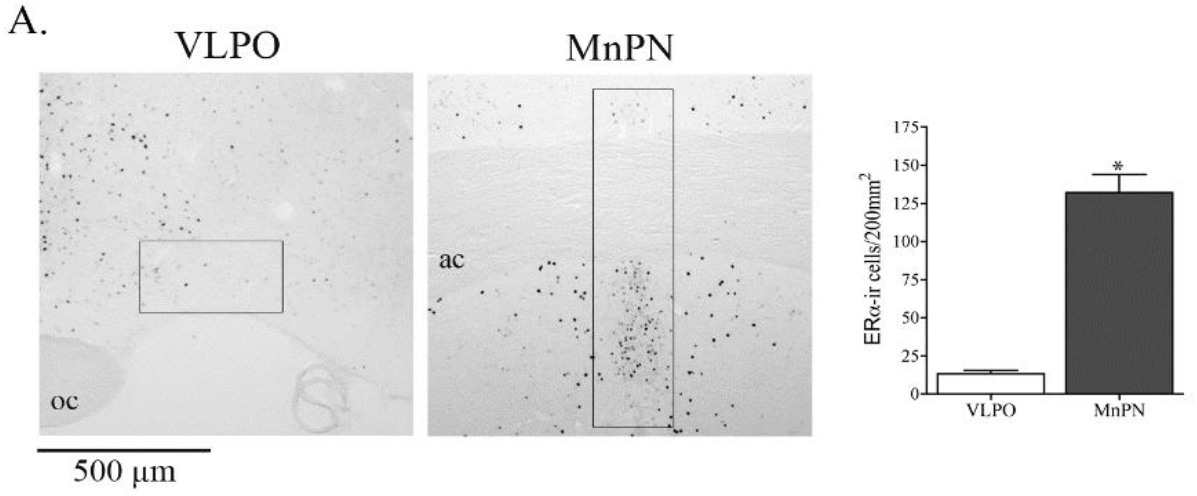
Estrogen Receptors are Highly Expressed in the Female MnPO. Staining for Estrogen Receptors (ER) show there is a high concentration in the MnPO but not the VLPO of females. (Student’s *t*-test, *, p<0.05 vs. VEH).

#### b. ER Antagonist Infusion to the MnPO Partially Rescues E2-Mediated Sleep Suppression

Building on the presence of E2-receptors in these nuclei, we attempted to test if the MnPO is necessary to mediate E2 actions on NREM sleep. Using the same exogenous E2 replacement paradigm shown to produce effects on sleep, we then cannulated the VLPO and the MnPO and infused ICI-182-780 (ICI), an estrogen receptor (ER) antagonist. This study attempts to determine if estrogen receptor (ER) signaling is required in either region for E2 suppression of sleep. These experiments show that blocking ER signaling at the MnPO, but NOT the VLPO, is able to ameliorate E2-mediated sleep suppression.

Due to greater expression of ER alpha in the MnPO, we ran a preliminary cohort of infusion of the direct estrogen receptor antagonist ICI 182,780 (ICI) into the MnPO. OVX rats were hormonally replaced with E2 or Oil, using the same paradigm as in Experiment 1, with a 5ug dose on Day 2 and a 10ug dose on Day 3. However, on days of E2 administration, an ER agonist, ICI, was also infused locally into the MnPO. (Fig. 10). Cannula targeting was confirmed by histology (Fig. 11).

**Fig. 10.**
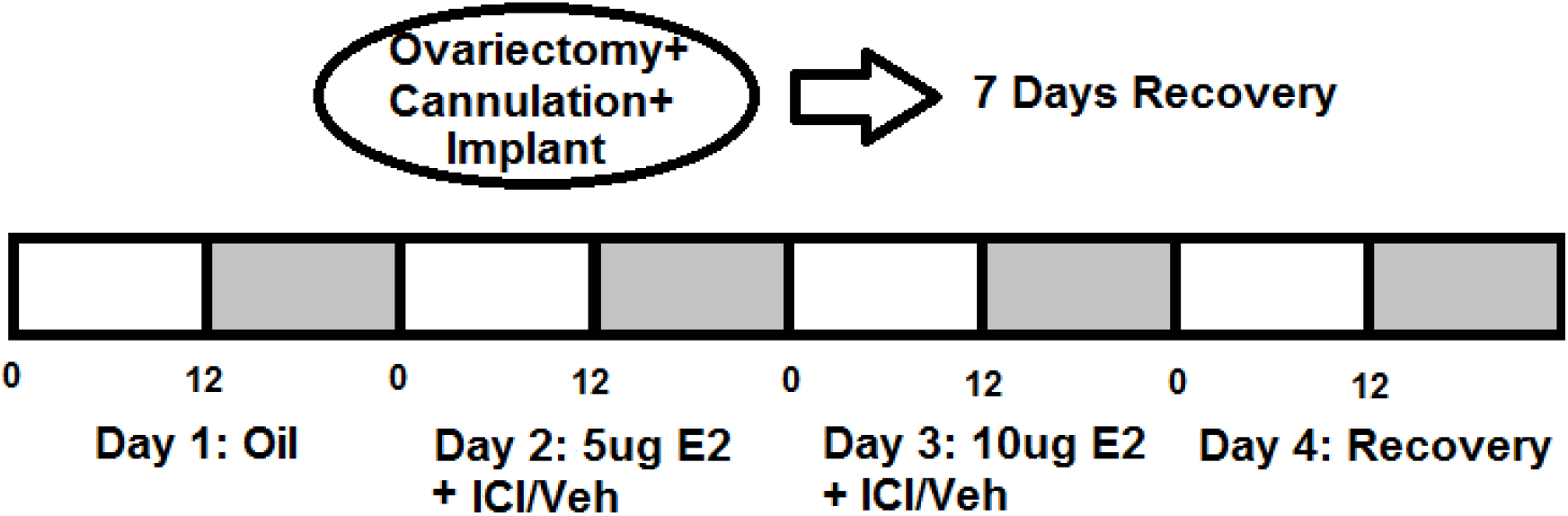
Timeline of ER Antagonism Experiment. Ovariectomized Sprague-Dawley rats (n=15) were treated with hormone replacement of either E2 or Oil using the same paradigm as in fig. 1–3 (5ug Day 1, 10ug Day 2) and cannulated to the MnPO. A subset of the animals (n=8) were treated as well with the Estrogen Receptor antagonist ICI through direct local infusion to the MnPO and the others (n=7) were treated with vehicle. Sleep behavior measured with EEG/EMG telemetry (DSI Inc., St. Paul MN).

**Fig. 11.**
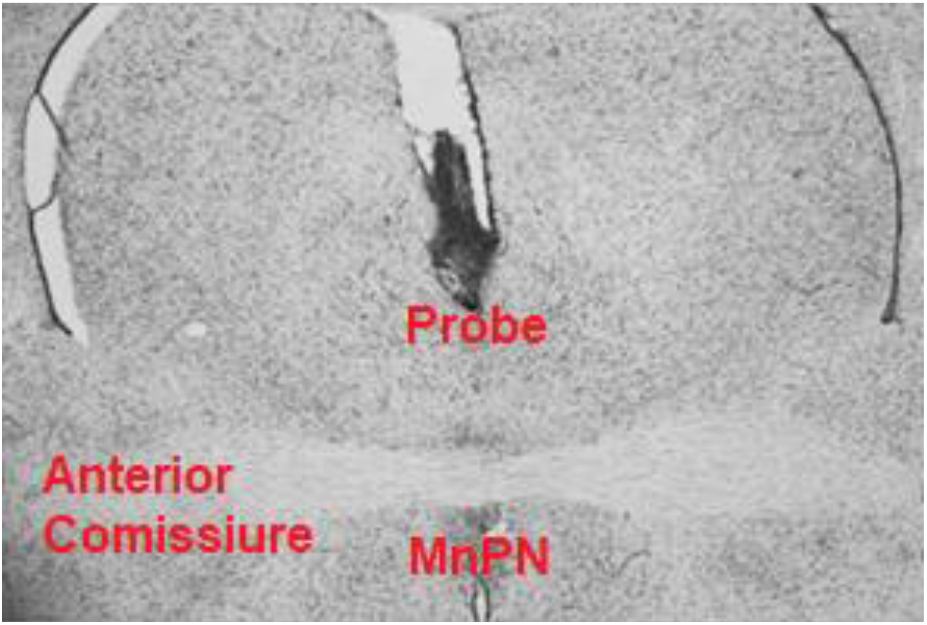
Representative Image of MnPO Probe Placement. Probe placement to the MnPO was confirmed histologically.

The effects of ICI in the MnPO on E2 suppression of sleep were moderate to high for wake *(d* = 0.67), total sleep *(d* = 0.67), NREM sleep (*d* = 0.5), and REM sleep (*d* = 1.36), prompting further investigation. In all groups, systemic E2 significantly modulated sleep-wake behavior; there was a main effect of E2 treatment for wake, total sleep, NREM sleep, and REM sleep during the dark phase. E2 treatment increased the time spent in wake and decreased sleep, both NREM sleep and REM sleep during the dark phase. In this study, pairwise comparisons of VEH infused animals given oil then E2 revealed that E2 treatment increased wake duration and decreased total sleep and REM sleep. Direct infusion of ICI into the MnPO significantly attenuated these effects during the dark phase, for both Wake and NREM sleep (Fig. 12A-B).

**Fig. 12.**
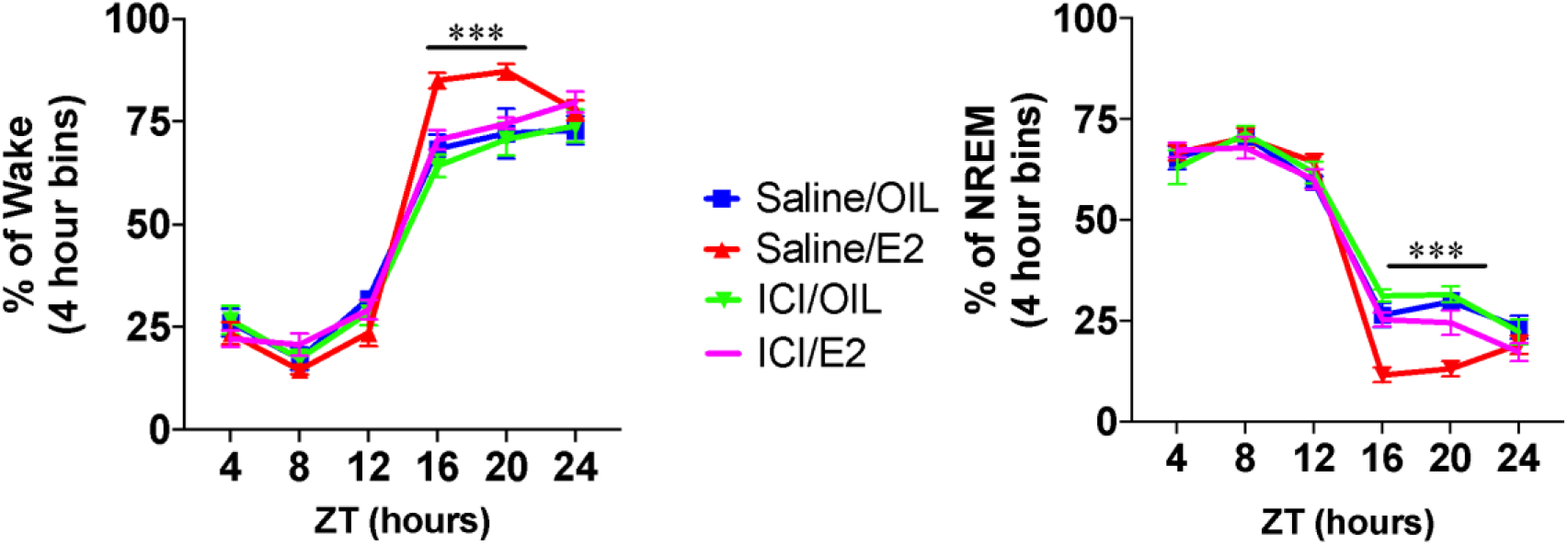
A-B. ER Antagonist ICI Reduces Wake Time and increases NREM Sleep Versus E2 Replacement. **(A)** Injection of ICI against a background of E2 treatment reduced wake time across much of the dark phase, with statistically significant decreases from ZT 12-16 (p<.001) and ZT16-20 (p<.001). There was no significant difference between ICI-treated and ICI-untreated animals without E2 replacement. **(B)** Injection of ICI against a background of E2 treatment increased NREM time across much of the dark phase, with statistically significant increases from ZT 12-16 (p<.001) and ZT16-20 (p<.001). There was no significant difference between ICI-treated and ICI-untreated animals without E2 replacement. Main effect of E2 treatment for wake, (F_1,12_ = 53.48, p < 0.001)/ (t_5_ = 2.56, p = 0.05), total sleep (F_1,12_ = 53.48, p < 0.001)/ (t_5_ = 2.68, p = 0.04), NREM sleep (F_1,12_ = 39.93, p < 0.001) and REM sleep (F_1,12_ = 57.03, p < 0.001)/ (t_5_ = 2.78, p = 0.04) during the dark phase.

Animals who received direct infusions of ICI into the MnPO exhibited about 47 minutes less wake time than VEH, about 37 more minutes of NREM sleep time, and 10 more minutes of REM sleep time (Fig. 13) during the dark phase. The percent change in wakefulness induced by E2 was not significantly different between VEH and ICI infusion groups. However, the percent changes in total sleep, NREM sleep, and REM sleep induced by E2 were significantly attenuated by ICI during the dark phase. As anticipated, the saline/E2 treated animals had significantly increased wake and reduced NREM sleep during the dark phase following the last injections. However, treatment with ICI (ICI/E2) blocked this E2 mediated effect, partially rescuing NREM and REM sleep, and inhibiting additional wake, to near baseline levels (Fig. 13). Keeping with the lack of significant effect of E2 in the light phase (Fig. 5), there was no effect of ICI on sleep times in the light phase (Fig. 14).

**Fig. 13.**
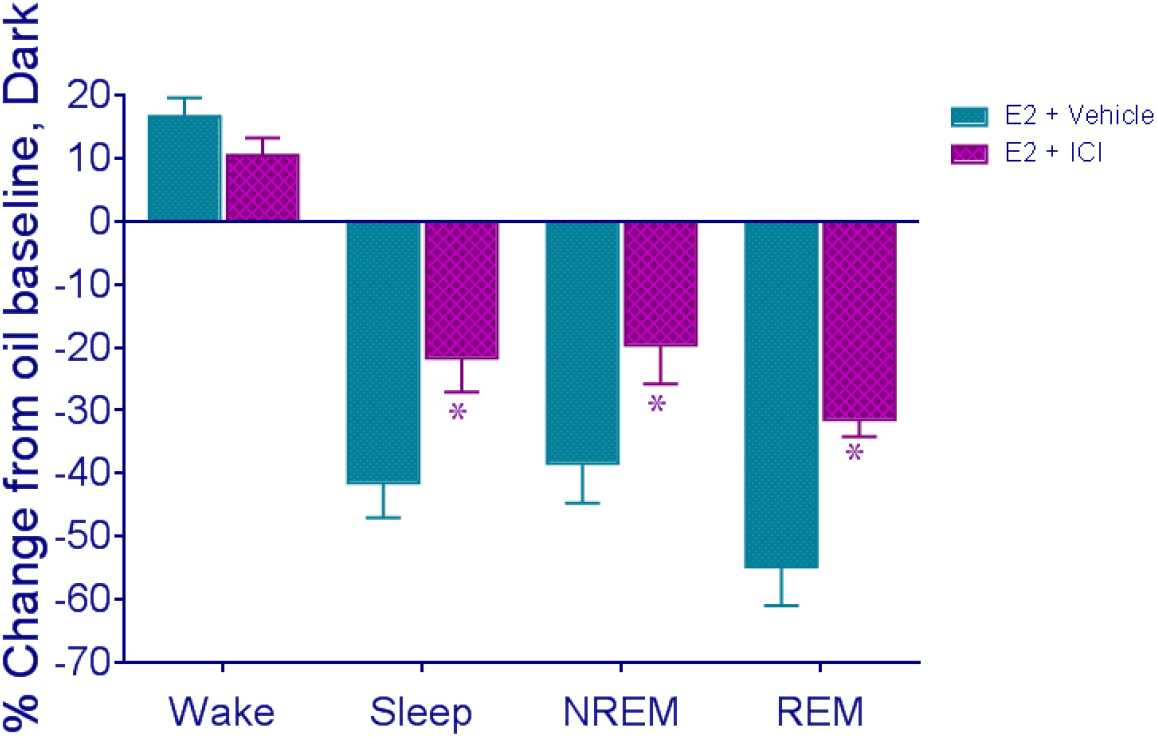
ICI Treatment Partially Rescues E2-*e* Mediated Dark Phase Sleep Suppression. We also compared total sleep time relative to each animal’s oil baseline recording, both with E2 treatment and vehicle and E2 and ICI treatment. During the Dark Phase, ICI treatment partially rescued the E2-mediated decrease in sleep, both in NREM and REM phases, leading to an increase in sleep time relative to E2+Vehicle animals. (Dark phase two-way ANOVA; main effect of. treatment, Wake: F(3,26) = 9.157; p<0.0005, NREM: F(3,26) = 14.86 p<0.0001) Wake (t_12_ = 2.376, *P* = 0.04) NREM sleep (t_12_ = 2.158, p = 0.05) REM sleep (t_12_ = 2.518, p = 0.03)

**Fig. 14.**
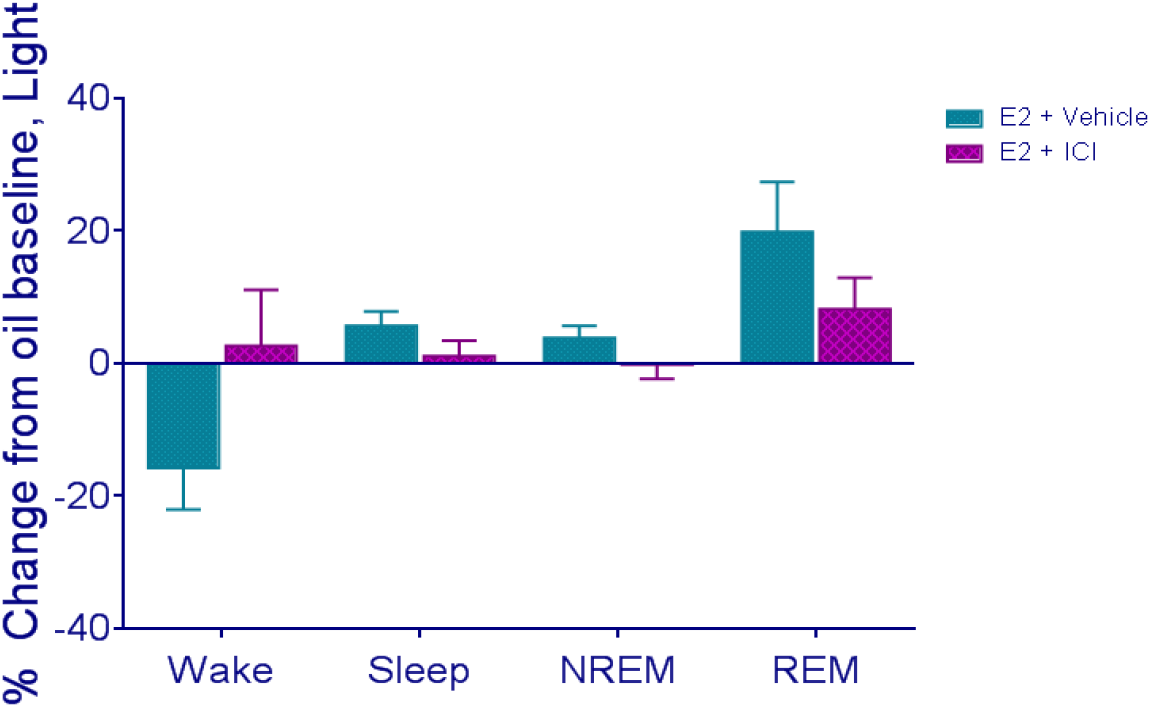
ICI does not Significantly Change Sleep in the Light Phase. As E2 did not have significant effects on sleep in the light phase relative to oil baseline, there was no effect of ICI antagonism of ERs in in the light phase relative to E2 treatment alone.

#### c. ER Antagonist Infusion to the VLPO Does NOT Rescue Sleep Behavior

Conversely to the MnPO, in the VLPO, infusions of ICI in the presence of systemic E2 had no effect on NREM or wake. Findings suggest that ICI infusion into the VLPO does not attenuate E2 effects on wake and sleep (Fig. 15A-B). Therefore, our data demonstrate that E2 acting directly in the MnPO and *not* the VLPO is necessary to attenuate NREM sleep, and that the inhibitory effect of E2 on sleep behavior is mediated by E2-expressing cells in the MnPO.

**Fig. 15.**
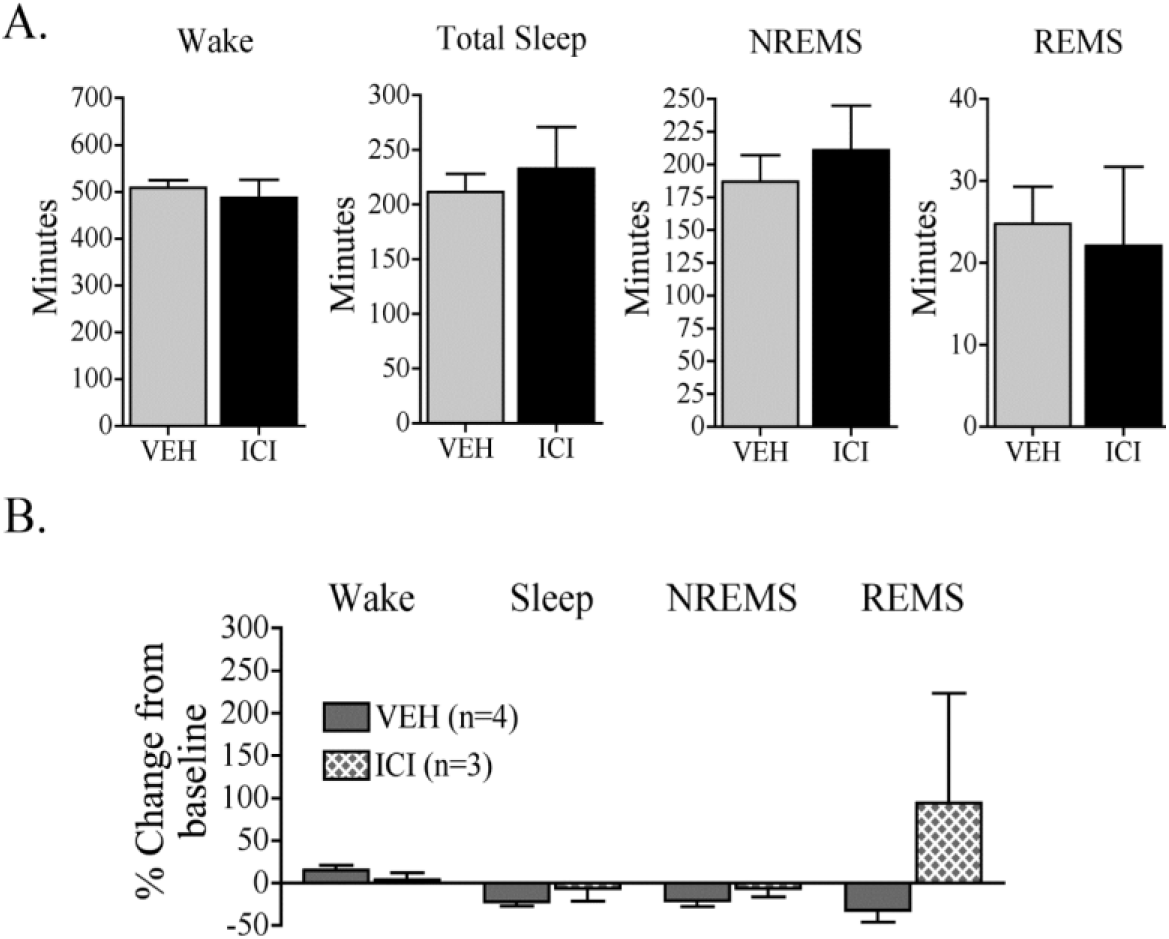
A-B. (Figure from DMC) ICI Infusion to the VLPO does not Change E2-I Driven Sleep Behavior. Unlike infusion into the MnPO, infusion of ICI into the VLPO did not affect wake time, total sleep time, or REM sleep time, either **(A)** in overall time or **(B)** change from oil control baseline. The effect size of the preliminary cohort for wake (*d* = 0.51), total sleep (*d* = 0.51), NREM sleep (*d* = 0.58), and REM sleep (*d* = 0.24) indicate any effect of ICI in the VLPO on E2 suppression of sleep was small to moderate.

### Experiment 3. Direct Infusion of Estradiol into the MnPO Increases Wake and Suppresses Sleep

Finally, we investigated whether E2 is sufficient to suppress sleep. To test this aspect of the signaling, we replaced the global subcutaneous administration of E2 with direct local infusion into the MnPO, to determine if E2 acting specifically at that nucleus is sufficient to reduce sleep. These studies address whether estrogen receptor signaling in the VLPO and/or the MnPO is the key site of action for the E2-mediated suppression of sleep in metrics of both necessity and sufficiency. OVX female rats were implanted with EEG/EMG telemeters and guide cannula to the MnPO. After recovery, animals were infused with 3 doses of cyclodextrin-encapsulated E2, a water-soluble form of E2, or equivalent amount of free cyclodextrin vehicle. This treatment was performed over 3 successive days at ZT18. (Fig. 16).

**Fig. 16.**
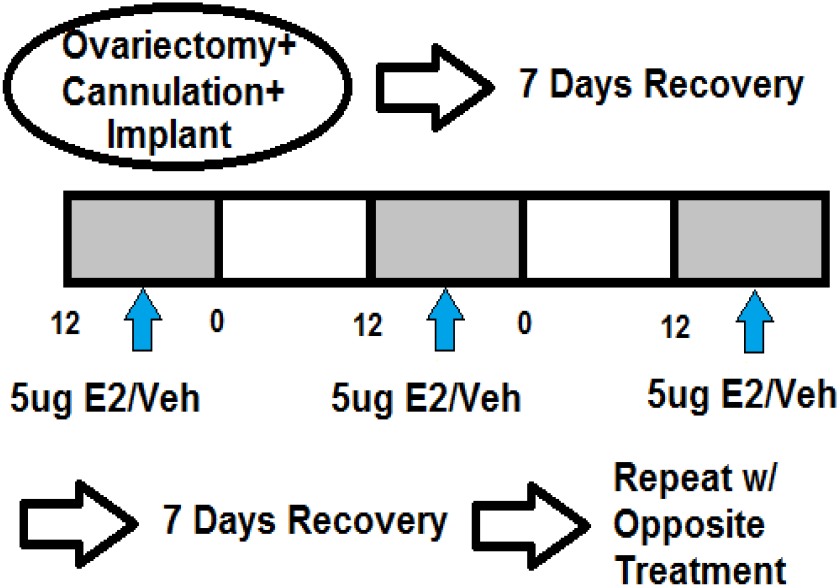
Timeline of Direct E2 Infusion Experiment. Female Sprague-Dawley rats (n=9) were ovariectomized and implanted with EEG/EMG telemeters and guide cannula to the MnPO. After recovery, animals were infused at ZT18 with 5ug cyclodextrin-encapsulated E2 in 5uL sterile saline, or 5ug cylodextrin vehicle in 5uL sterile saline. The same treatment was repeated for 3 successive days. After a 4-day washout, animals were subjected to the other treatment. Sleep architecture was quantified.

The significant differences in wake and NREM sleep were observed only in the light phase following the second injection. E2 infusion showed sleep suppression during the light phase on the second day of treatment, which significantly increased wake (Fig. 17) and decreased NREM sleep (Fig. 18) over the entirety of the light phase. There was no change in REM sleep (Fig. 19).

**Fig. 17.**
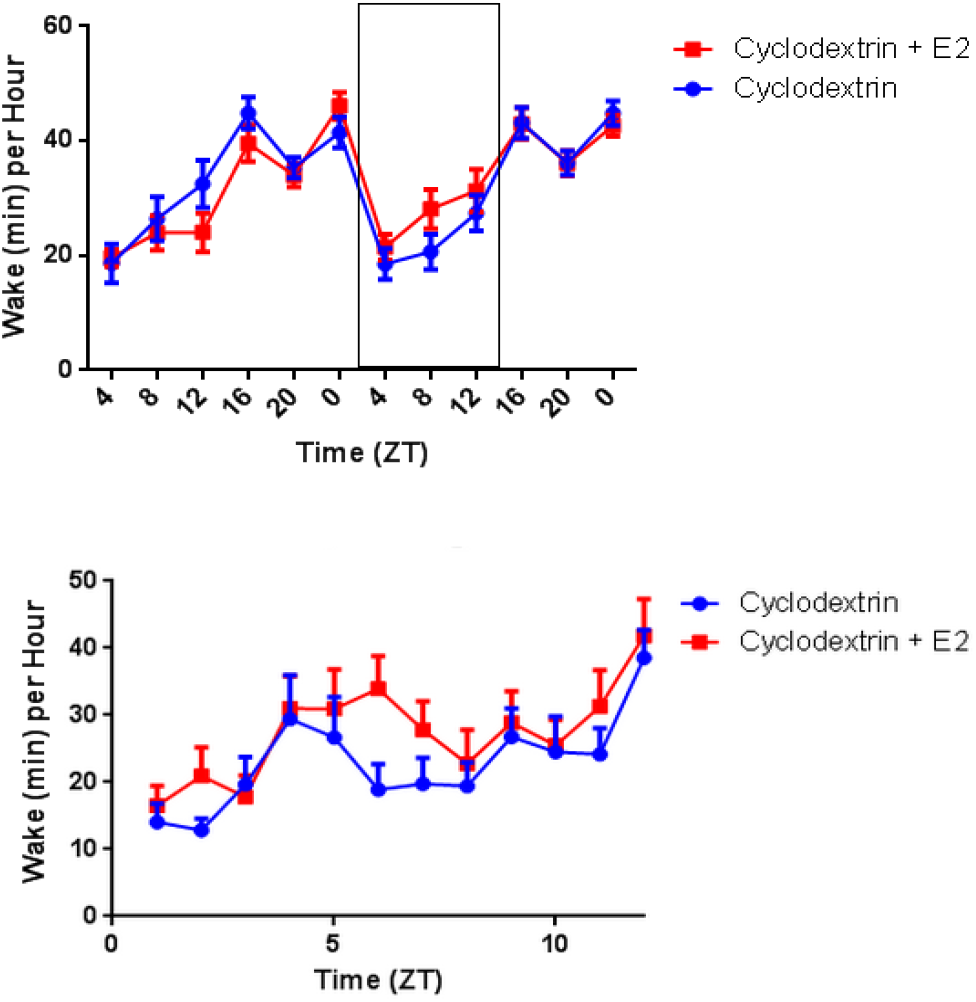
A-B. Direct E2 Infusion Increases Dark Phase Wake. E2-treated female rats showed an increase in wake time in the 2^nd^ light phase (p=.03, two-way ANOVA, main effect from ZT0-12 on treatment day 2). Lower Panel is detail of boxed region.

**Fig. 18.**
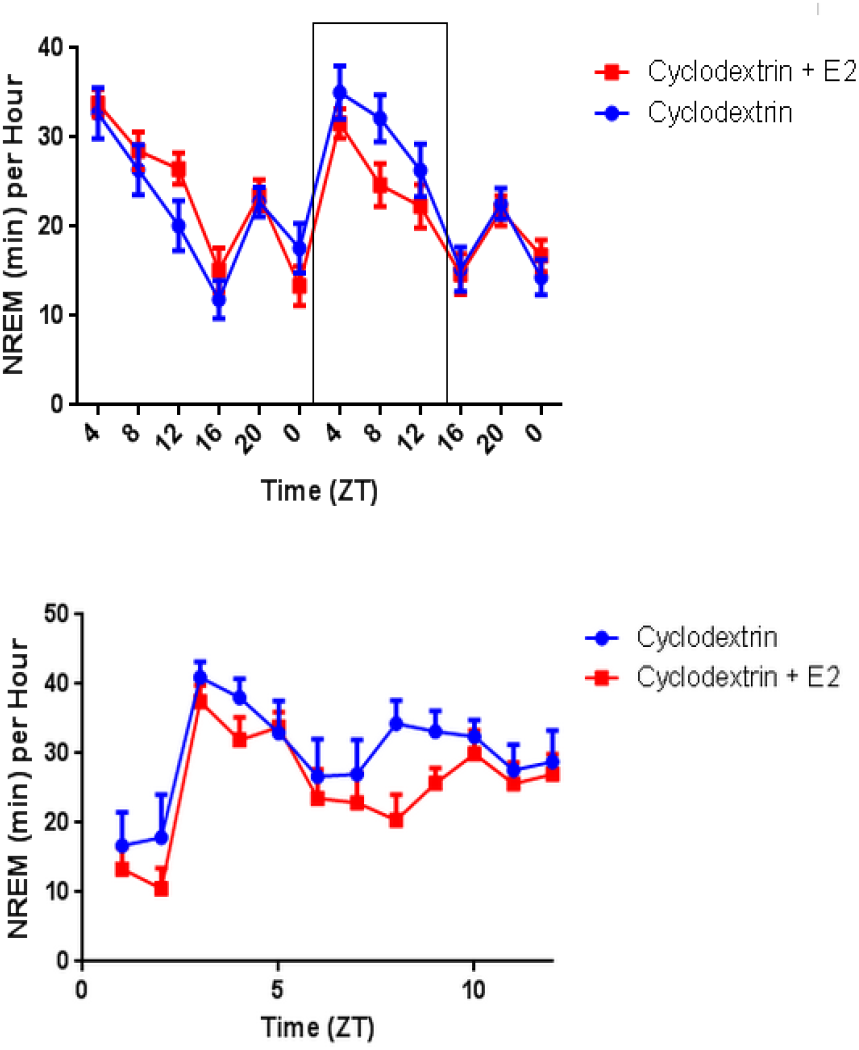
A-B.> Direct E2 Infusion Tends Toward a Decrease in Dark Phase NREM Sleep. There was no significant difference in NREM sleep time between the two groups, though the E2-treated female rats did show a trend (p=.10 main effect, two-way ANOVA ZT 0-12 day 2 of treatment) toward lower sleep time in the second light phase.

**Fig. 19.**
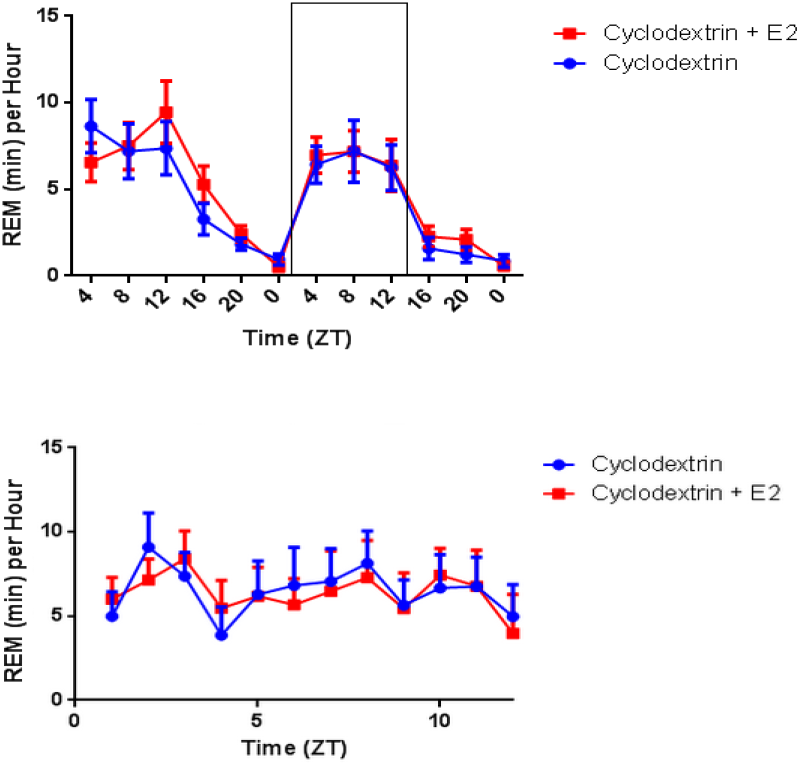
A-B. Direct E2 Infusion Has No Significant Effect on REM Sleep. There was no significant difference in REM sleep time between the two groups.

#### a. Estradiol Infusion Decreases NREM-SWA

We analyzed EEG spectral power in the second light phase with and without E2. While there was no significant change in NREM Delta Power over the entirety of the experiment, analysis of the second light phase showed decreases in delta power across many time points, particularly in the early part of the period (Fig. 20)

**Fig. 20.**
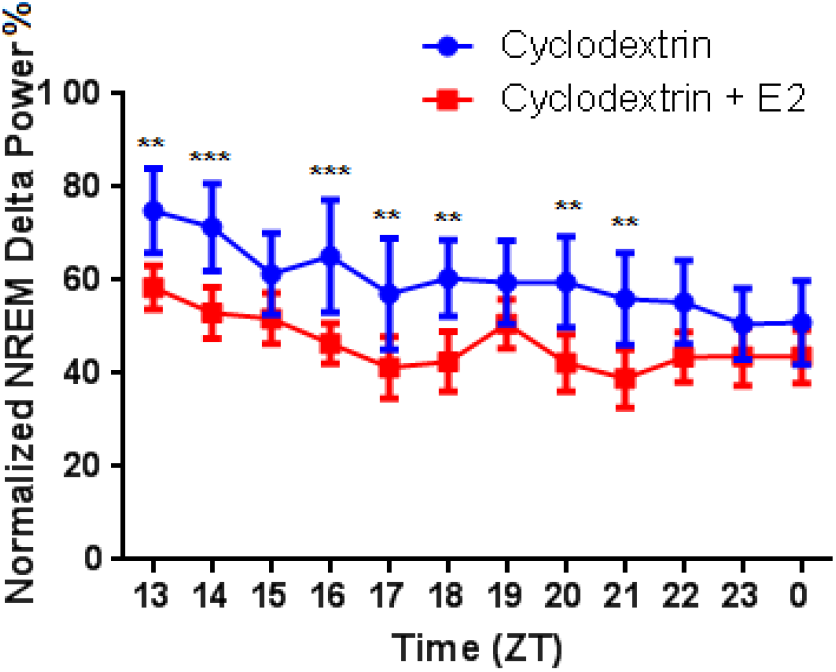
E2 Infusion Decreases NREM Delta Power. We analyzed the spectral power with and without E2. NREM Delta Power from E2-treated animals shows a significant decrease in several time points in the second light phase relative to oil controls. (Repeated Measures two-way ANOVA, main effect of hormone, F (1, 9) = 3.228P = 0.1059 post-hoc Sidak’s multiple comparison test, ZT 13 p<.01, ZT 14 p<.001, ZT 16 p<.001, ZT 17-18 p<.01, ZT 20-21 p<.01)

Further spectral analysis of this period shows that the effect is concentrated in the lowest frequencies of the delta band (below 2Hz), showing a decrease in the most coordinated brain waves that signify deep homeostatic sleep. (Fig. 21) Together, these findings strongly suggest that the MnPO is a direct mediator of E2 actions on sleep, and that E2 action at the MnPO is both necessary and sufficient for estrogenic effects on sleep.

**Fig. 21.**
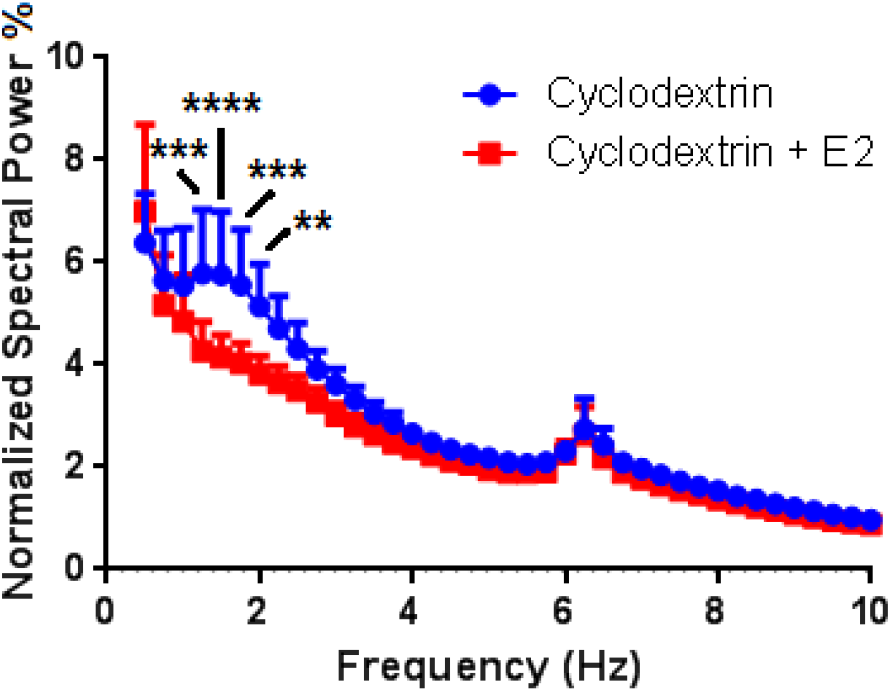
Direct Infusion of E2 Changes Power Most Significantly in the Low Delta Band. Spectral Fourier analysis revealed the decrease in delta power was localized to a particular portion of the low delta band, with the difference between E2 and vehicle significant at the 1.25 (p<.001), 1.5 (p<.0001), 1.75 (p<.001), and 2 (p<.01) Hz bands. (Repeated Measures two-way ANOVA, main effect of hormone, F (1, 9) = 4.129, post hoc Sidak’s multiple comparison test, 1.25Hz band p<.001, 1.5Hz band p<.0001, 1.75Hz band p<.001, 2Hz band p<.01).

## Discussion

Previous research studies using rodent models describe the changes in sleep across the female estrous cycle and following gonadectomy.^3, 6–7, 19^ Studies consistently reported that E2 suppresses NREM sleep and REM sleep in females, while changes in gonadal steroids cause little to no change in sleep in males. Here, we sought to address the mechanism by which proestrus levels of cycling ovarian steroids suppress sleep in females. We show that after hormone replacement of proestrus levels of E2, the suppression of sleep by endogenous hormones may be recapitulated. We further show that this suppression is due to the high levels of E2 alone, and that progesterone, the other major circulating ovarian steroid, did not have a significant impact on sleep behavior. Extending these findings, we found that E2 has direct actions within the sleep-active POA, specifically in the MnPO, which contains estrogen receptors (ERs). Antagonizing of ERs in the MnPO, but not the VLPO, attenuated the E2-mediated suppression of both NREM and REM sleep. We finally found that, in addition to E2 actions at the MnPO being necessary for E2 suppression of sleep, it is also sufficient, as the direct infusion of E2 into the MnPO suppressed sleep with no other intervention. Based on our findings, we predict that proestrus levels of E2 alone, acting at the MnPO, mediate sex-hormone driven suppression of sleep in female rats.

From our findings, we further conclude that E2 is both necessary and sufficient to reduce the activation of MnPO sleep active cells, thereby releasing the inhibitory tone on downstream targets. The MnPO contains GABAergic sleep-active projection neurons, which innervate the lateral hypothalamus and multiple brainstem nuclei.^16^ GABAergic MnPO neurons have direct inhibitory control over the orexinergic neurons in the perifornical area/ lateral hypothalamus.^167^ These orexinergic neurons are a key source of arousal signaling, suggesting a sleep-promoting mechanism of the MnPO. Since ICI had little to no effect within the VLPO, while E2 in the MnPO was sufficient to induce changes, E2 is most likely acting predominantly on the MnPO and *not* acting directly on the neural circuits of the VLPO. However, as the MnPO also innervates the VLPO,^31^ a decrease in MnPO activation by E2 may elicit a similar decrease downstream in the VLPO.

## Estradiol Acts at the MnPO to Affect Sleep Times

The molecular mechanisms and site of action of hormonal impacts on sleep are poorly understood, and studies investigating where and how female steroids act on the brain are only an emerging area of investigation. From our findings, we predict that E2 is both necessary and sufficient to reduce the activation of MnPO sleep active cells, and release the MnPO-driven inhibitory tone on downstream targets; however, the pathway of E2 action on sleep is likely far broader. Steroid receptors, particularly for estrogen, are present throughout the brain and prevalent on multiple sleep-regulating nuclei such as the basal forebrain.^32^ Downstream, the orexinergic wake-promoting system of the lateral hypothalamus receives inputs from the MnPO and is highly sensitive to fluctuations in endogenous and exogenous ovarian steroids,^2^ suggesting that this section of the homeostatic sleep/wake circuitry may be a key site for estrogen action.

Since ICI had little to no effect within the VLPO, while E2 in the MnPO was sufficient to induce changes, E2 is most likely acting predominantly on the MnPO and *not* acting directly on the neural circuits of the VLPO; however, indirect actions on the VLPO via MnPO neurons seem likely to be present. The decrease in Fos-ir cells in the presence of E2 may be the result of reduced inhibition. Thus, an E2-driven decrease in MnPO activation may lead to a downstream decrease in the VLPO. Additionally, the sex difference in ER alpha expression in the MnPO may account for the difference in sensitivity of males and females to the suppressive effects of E2 on sleep.^7^ Further experiments are necessary to determine how ERs may be activated in the MnPO, and any downstream effects triggered, through our necessary-and-sufficient E2 dosing paradigm.

The multiple potential sites of E2 action present an interesting question of how E2 is working to disconnect molecular measures of sleep need from sleep behavior thought to be characteristically directly responsive to that need. We acknowledge that these experiments are heavily focused on the MnPO as one potential key site and that more global actions of E2 within other sleep and arousal circuits may also be important. However, given that we understand so little about how E2 is influencing sleep, expanding these studies to other brain regions and molecular mechanisms in the future could provide more insight into how E2 and sleep networks interact to affect sleep.

## Other Potential Sites and Mechanisms Exist for Estradiol Suppression of Sleep

Sleep-wake behaviors are regulated by reciprocal connections between sleep-promoting nuclei in the preoptic area (POA) and arousal centers in the hypothalamus and brainstem.^14^ The ventrolateral preoptic area (VLPO) and the median preoptic nucleus (MnPO) are two key sleep-active nuclei involved in the onset and maintenance of sleep.

Both regions express Fos, a proxy marker for neuronal activation, during sleep periods, which correlates inversely to the amount of previous sleep.^9, 14, 33–34^ Thus, the neuronal activation of these regions can be seen as a proxy for sleep pressure, or homeostatic sleep need. Moreover, the VLPO has long been shown to have sensitivity to changes in ovarian steroids. E2 replacement following ovariectomy reduces both Fos expression and protein expression of lipocalin-type prostaglandin D synthase (L-PGDS), the synthesizing enzyme for the somnogen prostaglandin D2, within the VLPO of female rats compared to ovariectomized controls.^19^ These POA nuclei are also proposed to modulate homeostatic sleep drive. Together, these data suggest that E2 alters critical factors involved in sleep, particularly in the sleep-active POA.

Our findings are the first report of direct actions of E2 on sleep behavior within a sleep-active nucleus. It is clear, however, that the MnPO is only one region mediating E2 suppression of sleep. Local ER antagonism in the MnPO only partially rescued baseline sleep and wake. Therefore, it is likely that E2 has direct actions in other nuclei regulating sleep/wake. For example, the histaminergic neurons in the tuberomammillary nucleus may be targets for E2. Histamine is involved in arousal and is a direct downstream target of the VLPO.^33–35^ Fos expression is higher in ovariectomized females treated with E2 compared to oil, indicating increases in arousal following E2 treatment.^19^ It is unclear if this is a direct or indirect effect of E2. Nuclei within the ascending reticular activating system contain ERs and are potential target sites for E2-mediated arousal.^36–41^ The noradrenergic neurons of the locus coeruleus (LC) are involved in arousal and contain binding sites for E2, suggesting that E2 may directly increase arousal by increasing activation of the LC. Additionally, the cholinergic neurons of the basal forebrain and pedunculopontine tegmental nucleus/laterodorsal tegmental nucleus, which play a key role in cortical activation during arousal, can also be target sites for E2.^42^ Increased activation of these regions by E2 may account for increased high frequency oscillations in sleep EEGs, which may be an objective measure of poor sleep quality.^43^

Alternatively, E2 may act within other sleep-promoting nuclei like those involved in REM sleep generation. It is possible that E2 either directly or indirectly reinforces the activity of REM-OFF cells in the brainstem.^44^ MCH/GABAergic neurons are proposed to control the switch into REM sleep by inhibiting REM-OFF cells.^45^ MCH neurons do not express ER;^46^ however, the neighboring GABAergic population in the lateral hypothalamus may do so. E2 may reduce the activity of MCH/GABAergic cells (directly or indirectly) to suppress REM sleep.

## Potential Molecular and Neurological System Mechanisms of E2 Effects on Sleep

Beyond the question of a site of action, the question of how, in terms of molecular and neurological mechanism, E2 may be meditating sleep effects is an important one. Two distinct systems govern aspects of sleep regulation, the circadian and homeostatic sleep drives, which operate in concert to generate an overall sleep/wake cycle that is responsive to both the animal’s intrinsic homeostatic needs as well as external factors such as the light-dark cycle. The homeostatic drive, which governs the amount of sleep needed after a given period of wake to maintain homeostasis, independent of circadian factors, is thought to utilize both the VLPO^8^ and MnPO^2^ as key originators of this pathway. The VLPO and MnPO send GABAergic projections to key mediators of the wake state, including nuclei in the lateral hypothalamus governing the orexinergic wake system.^2^ Additionally, the VLPO and MnPO have been shown as sites of sensitivity to adenosine, an important mediator of homeostatic sleep pressure.^47^ Further exploration of these molecular and neurological pathways could provide greater insight into precisely how E2 is affecting sleep need and behavior.

## Conclusion

Rodents provide a model system for studying the mechanism underlying the sensitivity of the sleep circuitry and behavior to E2. Such a model is highly significant in the identification of neuronal targets for E2 within the sleep circuitry. Here, we describe the key role of E2 alone in modulating sleep behavior, as well as provide first clear evidence of a direct role for E2 in a sleep-active nucleus. The identification of the MnPO as a direct site of E2 action, showing that it is both necessary AND sufficient for induction of estrogenic effects on sleep, now allows for more mechanistic research to determine how E2 is suppressing sleep in females. Understanding the circuits that E2 can act on to regulate sleep may enable better drug development and treatment of sleep disorders in the clinical population.

